# Value control of punishment

**DOI:** 10.64898/2026.02.22.707274

**Authors:** Felipe I. Varas, Anthony Dickinson, Omar David Perez

**Author notes:** **Author Note** Correspondence regarding this article should be sent to Omar David Perez or Felipe I. Varas. **Authors’ contributions** ODP and AD designed the study. FIV programmed the tasks and ran the experiment. FIV analyzed the data. ODP, FIV and AD drafted the manuscript. ODP acquired funding. All authors read and approved the final manuscript.

## Abstract

Punishment raises a basic question for human cognition: do agents suppress actions because they represent the negative consequences those actions produce? Everyday deterrence, from parental reprimands to legal sanctions, assumes that people regulate behavior by learning what their actions cause and by evaluating those consequences negatively. However, in spite of this widely-held assumption, experimental evidence that punished instrumental behavior is controlled by the current value of the punisher is currently lacking. To test this idea, we applied an outcome revaluation procedure to instrumental punishment in rats. Rats first learned to press two levers for food. One lever was then additionally punished by response-contingent footshock, whereas the other remained unpunished. The aversiveness of the shock was subsequently reduced by pairing it with food in the absence of the levers. To test if the punisher’s value was part of the animals’ representation of the consequences of their actions, responding was tested in extinction, with no shock delivered. We found that revaluation of the shock selectively abolished the suppression of the punished response. Thus, punishment was not a fixed behavior controlled by the prior occurrence of shock, but depended on the shock’s current value. Our results show that instrumental punishment can be controlled by a value-sensitive representation of the aversive consequence produced by the action and is, in this sense, structured in the same manner as canonical goal-directed reward seeking.

## Introduction

A central assumption in human cognition is that agents regulate their actions according to their consequences. When an action is followed by an aversive consequence, that action tends to be suppressed. A child may stop touching a heater after a burned finger, or stop throwing tantrums if these are followed by parental reprimands. Likewise, adults may refrain from speeding or evading fares because they expect these actions to produce fines or sanctions. Punishment is also central to human institutions that aim to regulate behavior. Deterrence policies, for example, assume that legal penalties reduce unwanted actions because people learn what their actions cause and evaluate those consequences negatively (Becker, 1968).

These examples all share a common assumption: agents refrain from some behaviors because they learn that the action causes a punishing outcome which, being experienced or represented as aversive, costly, painful, or otherwise undesirable, is assigned a negative value. The interaction between this instrumental knowledge and outcome evaluation then leads the agent to refrain from the action. However, in spite of its common-sense appeal, there is so far no direct behavioral evidence that punished behavior is controlled by the current value of the punisher.

This issue is important because psychological approaches from animal learning theory explain suppression after punishment from a different perspective. The punished response may become inhibited, the context may acquire suppressive properties, or the aversive event may generate a general emotional state that reduces behavior. In all these cases, punishment would suppress responding without requiring that performance be guided by a representation of the specific action–punisher relation and the punisher’s current value. The Conditioned Emotional Response account, for example, explains punishment through a Pavlovian emotional reaction that competes with instrumental performance (Estes, 1944) and the negative Law of Effect treats punishment as weakening the tendency to perform the punished response (Thorndike, 1911). These accounts differ in important ways, but they share a common feature: they explain suppression without requiring that behavior be guided by a representation of the punisher’s current value.

Punishment, however, can be sensitive to the contingency between a response and its punisher. Bolles and colleagues, for example, trained rats to perform two topographically distinct responses for food and then punished one of them (Bolles et al., 1980). Although both responses were suppressed early in punishment training, suppression later became specific to the punished response. Jean-Richard-dit-Bressel and McNally (2015) replicated this response-specific suppression using two retractable levers, and Broomer and Bouton (2023) found response-specific punishment in a discriminated instrumental procedure. Therefore, punishment is not simply facilitating an automatic, incompatible response.

However, response specificity alone is not sufficient to conclude that the response is outcome-value driven. In both humans and non-human animals, value-driven performance requires not only that the subject tracks which response was made in relation to the punisher, but also the current value of the outcome that response produces (de Wit & Dickinson, 2009; Dickinson et al., 2018; Perez & Dickinson, 2020).

This criterion is well established in the study of rewarded instrumental actions. To test if a response is sensitive to outcome value, an animal first learns that a response produces a rewarding outcome, but then the response is withheld and the outcome is independently revalued. Finally, the response is tested in extinction, when the revalued outcome is not available. Because the outcome is absent at test, any changes in responding can only reflect a prior representation of response-outcome learning and the outcome’s current value. In other words, the subject knows that certain actions are followed by a specific outcome, the value of that outcome, and responds according to that causal representation, not because they re-experience the outcome in its new value. Therefore, if performance changes appropriately after revaluation with respect to the outcome’s new value, the action is considered value-driven (or goal-directed). Adams and Dickinson (1981) first used this assay to show that rat lever pressing for food can be sensitive to value, and subsequent work has extended this framework to human instrumental action (de Wit & Dickinson, 2009; Dolan & Dayan, 2013).

Unlike reward learning, showing that instrumental punishment is value-driven is significantly more challenging. The revaluation procedure, in this case, should be applied to the punisher, which has negative incentive value. Therefore, in order to establish whether punishment is value-driven with respect to the punisher, one must seek to reduce the aversive value of the punisher and ask whether this change affects performance of the punished action in extinction, without further experiences with the revalued punisher. For example, a person who usually evades bus fares should stop evading if she is informed that the fines have become harsher, without having to experience being caught (and fined) with this higher cost.

Previous studies have suggested that it is possible to reduce the aversive value of a punisher. Pearce and Dickinson (1975) showed that shock-food counterconditioning, which requires systematically pairing the shock with a rewarding outcome, reduces the general aversiveness of shock in conditioned emotional response procedures, and Dearing and Dickinson (1979) showed that making shock a signal for water reduced its capacity to punish lever pressing. Although these studies suggest that the value of a punisher can indeed be altered by conditioning procedures, they do not show whether punished actions are controlled by the current value of the punisher, which requires revaluing the outcome independently and testing whether it can change punished responses in an extinction test.

To test if punishment is controlled by outcome value, we applied the outcome revaluation logic to instrumental punishment. Rats first learned to press two levers concurrently for food. Then, during punishment, one lever was still rewarded but also delivered response-contingent footshock, while the other remained rewarded but unpunished. The aversiveness of the shock was then reduced by pairing it with a rewarding outcome in the absence of the levers; in the control group, food was paired with a neutral cue. We then tested responding on both levers in extinction, with no shock delivered. If punishment is controlled by a causal representation of the response-shock relation and the current value of the shock, then revaluing the shock should selectively restore responding on the punished lever. To our knowledge, this is the first direct test of whether behavior under punishment is controlled by the current value of a primary biological punisher.

## Methods

### Subjects

Forty-eight female Sprague-Dawley rats from the biotherium of the Faculty of Chemical and Pharmaceutical Sciences at Universidad de Chile were run in two batches of 24. Batch was included as a between-subjects factor in all analyses. The sample size (N=48) was determined based on achieving sufficient power to detect a strong effect size (d > 0.6) typical in outcome revaluation studies with similar designs.

Rats began the experiment at approximately 9 weeks of age (*M* = 234 g, range: 202–271 g). Subjects were pair-housed under a 12:12-h light/dark cycle with all sessions conducted during the dark phase. They had ad libitum water and were food-restricted to 85% of free-feeding weight. All procedures were approved by the Comité Institucional de Cuidado y Uso de Animales (CICUA) at Universidad de Chile (Protocol #2508-FCS-UCH).

### Apparatus

Eight identical operant conditioning chambers (ENV-008; 29.53 × 24.84 × 18.67 cm, L x W x H), manufactured by Med Associates, were housed in sound-attenuating cubicles. Each contained two retractable levers (ENV-112CM; 4.8 cm long, protruded 1.9 cm when extended), a recessed food magazine (ENV-202RMA; 5.1 × 5.1 cm) between them, a houselight (ENV-215M-LED), and a grid floor (ENV-005; nineteen stainless steel rods, 0.47 cm diameter, spaced 1.57 cm apart) connected to a scrambler (ENV-414) for delivering footshock. Magazine head entries were recorded continuously via an infrared detector (ENV-254-CB). Food reinforcers consisted of 45-mg sugar pellets (locally manufactured).

### Procedure

Table 1 provides an overview of the experimental design. Briefly, rats first acquired lever pressing for food reinforcement, then received punishment on one lever via response-contingent footshock. Following punishment, the aversive value of the shock was manipulated through counterconditioning before a final revaluation test assessed the effect of this manipulation on punished responding. Each phase is described in detail below.

**Table 1.**
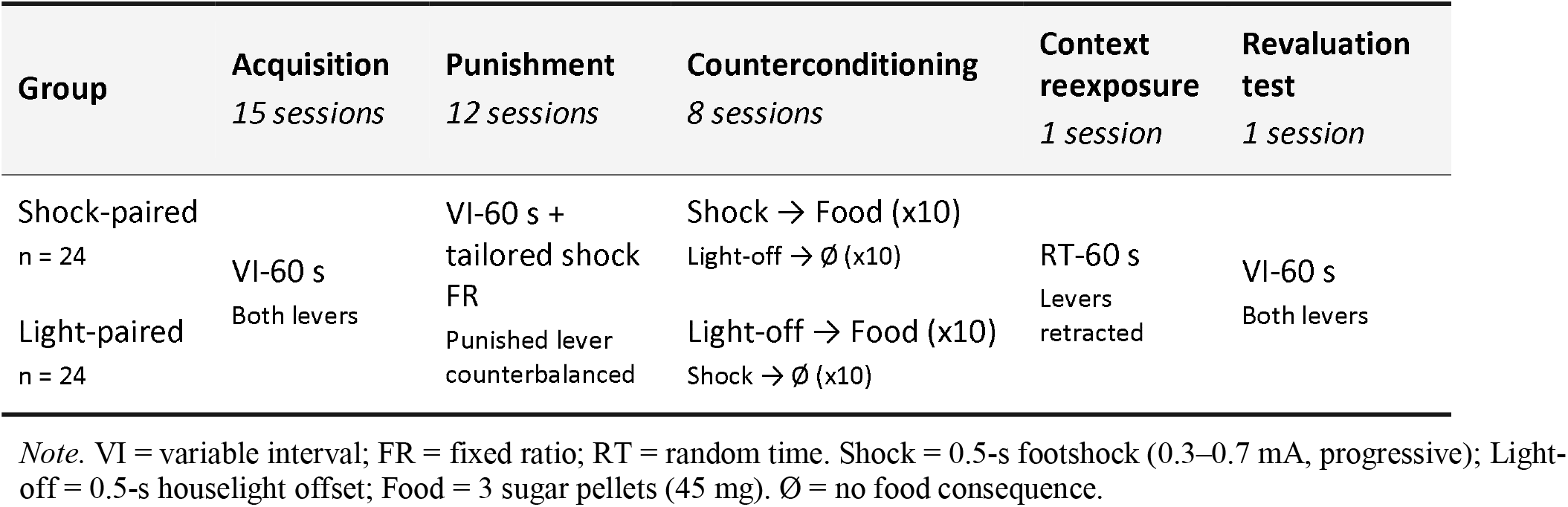
Overview of experimental procedure.

#### Pretraining

Four days of pretraining shaped lever pressing from magazine training (RT 60-s) through FR-1 (Days 2–3) and FR-5 (Day 4). All sessions lasted 30 min. From Days 2–4, one lever was available for the first 15 min, after which it retracted and the second lever was extended for the remaining 15 min. The order of lever presentation was counterbalanced across subjects.

#### Acquisition

Each rat received 15 daily 30-min sessions in which both levers were available concurrently, each reinforced on its own independent schedule. Reinforcers were delivered on an independent variable-interval (VI) 60-s schedule for presses on each lever with a 2-s changeover delay. Intervals were sampled from an exponential distribution (mean = 60 s, maximum = 180 s).

#### Punishment

Each rat received 12 daily 30-min sessions in which the VI 60-s schedules continued on each lever independently, with each lever programmed on its own schedule regardless of responding on the other. Response-contingent footshock was superimposed on one lever (counterbalanced) according to an individually tailored fixed-ratio schedule, calculated so that the subject would receive approximately 10 shocks per session at its baseline response rate. Shock intensity increased progressively from 0.3 mA (0.5 s) to a terminal intensity of 0.7 mA (0.5 s). This gradual escalation was designed to minimize the development of competing freezing responses that would interfere with lever pressing, following the procedural recommendations of Fernando et al. (2015).

#### Counterconditioning

The levers were withdrawn and rats were randomly assigned to two groups (*n* = 24). In the shock-paired group, each footshock was followed by three sugar pellets while a 0.5-s houselight offset was presented without food. The light-paired group received the reverse arrangement. Each of 8 daily 30-min sessions contained 10 shock and 10 light-off presentations at pseudo-random intervals (*M* = 82.74 s). This design ensured that both groups received identical numbers of shock presentations and food deliveries, controlling for the effects of mere shock exposure, shock habituation, and differential experience with the appetitive reinforcer.

#### Context reexposure

One day prior to the revaluation test, all rats received a single 30-min session in the same operant chambers used throughout training, with levers retracted and food delivered on an RT 60-s schedule in the absence of shock. This session was designed to minimize potential group differences in contextual conditioning and to re-establish an appetitive context.

#### Revaluation test

All rats received a 30-min test with both levers reinserted and reinforced on the concurrent VI 60-s schedules but in the absence of any shocks.

### Data analysis

All statistical analyses were conducted using R (version 4.5.1). Lever pressing (responses per minute, rpm) was the dependent variable for acquisition, punishment, and revaluation test analyses. Responding during the final acquisition session was compared across groups using a between-subjects ANOVA with Group (Shock-paired vs. Light-paired), Lever (control vs. punished), and Batch as factors. Punishment was evaluated using a mixed ANOVA with Lever and Session (Final acquisition session + 12 punishment sessions) as within-subjects factors, and Group and Batch as between-subjects factors. For the revaluation test, a mixed ANOVA was conducted with Lever and Phase (Baseline: final punishment session; Test: Revaluation test) as within-subjects factors, and Group as between-subjects factors. Including the final punishment session as a baseline phase controls for pre-existing individual differences in punishment-induced suppression and provides a more sensitive test of the hypothesis that counterconditioning selectively reinstated punished responding—a prediction that concerns change from a pre-manipulation baseline rather than absolute responding at test. To quantify selective recovery at the individual level, a difference-in-differences recovery score was computed as (Punished Test - Punished Baseline) (Control Test - Control Baseline). To examine the temporal dynamics of lever pressing within the revaluation test, responding was additionally modeled across 5-min time bins using a linear mixed-effects model fitted with the *lme4* package (Bates et al., 2015), with Group, Lever, their interaction, and a linear time term as fixed effects, and by-subject random intercepts and random slopes for Lever and time with freely estimated correlations: (1 + Lever + bin | Subject), following the recommendation to use the maximal random effects structure justified by the design (Barr et al., 2013). Model selection was based on AIC comparison across six candidate models; parameters were estimated using REML with the bobyqa optimizer, fixed effects significance was evaluated using Satterthwaite’s approximation as implemented in *lmerTest* (Kuznetsova et al., 2017). Counterconditioning was assessed using a mixed ANOVA on head entry difference scores (post-stimulus minus pre-stimulus within 4-second windows) with Signal (Paired vs. Unpaired) and Session (8 counterconditioning sessions) as within-subjects factors, and Group and Batch as between-subjects factors. To assess potential group differences in contextual conditioning prior to the revaluation test, head entries were also analyzed during a context reexposure session (RT 60-s) in which no levers were present. Head entry rates were aggregated into 3-min time bins across the 30-min session and submitted to a mixed ANOVA with Bin (10 levels) as a within-subjects factor and Group and Batch as between-subjects factors. Greenhouse-Geisser corrections were applied when sphericity assumptions were violated. Planned comparisons were conducted, as they were derived directly from the a priori theoretical prediction that counterconditioning would selectively reinstate punished responding in the shock-paired but not the light-paired group (Rosenthal & Rosnow, 1985); post-hoc comparisons, including those for the context reexposure analysis for which no directional predictions were made, used Bonferroni correction. Effect sizes are reported as generalized eta-squared (*ges*) for omnibus ANOVA effects and Cohen’s *d* for planned comparisons. For visualizations of within-subjects designs, error bars represent within-subjects standard errors corrected using the Cousineau-Morey method. Statistical significance was set at α = .05.

## Results

The rate of lever pressing increased progressively across acquisition sessions. On the final session (Session 15), mean response rates (SD) were as follows: Shock-paired group: control lever = 10.4 rpm (5.0), punished lever = 10.8 rpm (5.5); Light-paired group: control lever = 10.7 rpm (7.09), punished lever = 11.3rpm (6.0). A Group x Lever x Batch ANOVA yielded no significant main effects nor interactions, all *ps* > .15.

### Punishment

A mixed ANOVA with Condition and Session as within-subjects factors, and Group and Batch as between-subjects factors confirmed successful punishment learning. There was a significant main effect of Condition, F(1,44) = 26.46, *p* < .001, *ges* = .173, indicating greater responding on the control lever than the punished lever. The Condition x Session interaction was also significant, F(12,528) = 22.03, *p* < .001, *ges* = .058 (Greenhouse-Geisser corrected, ε = .247), reflecting progressive suppression on the punished lever across sessions (see Figure 1).

**Figure 1.**
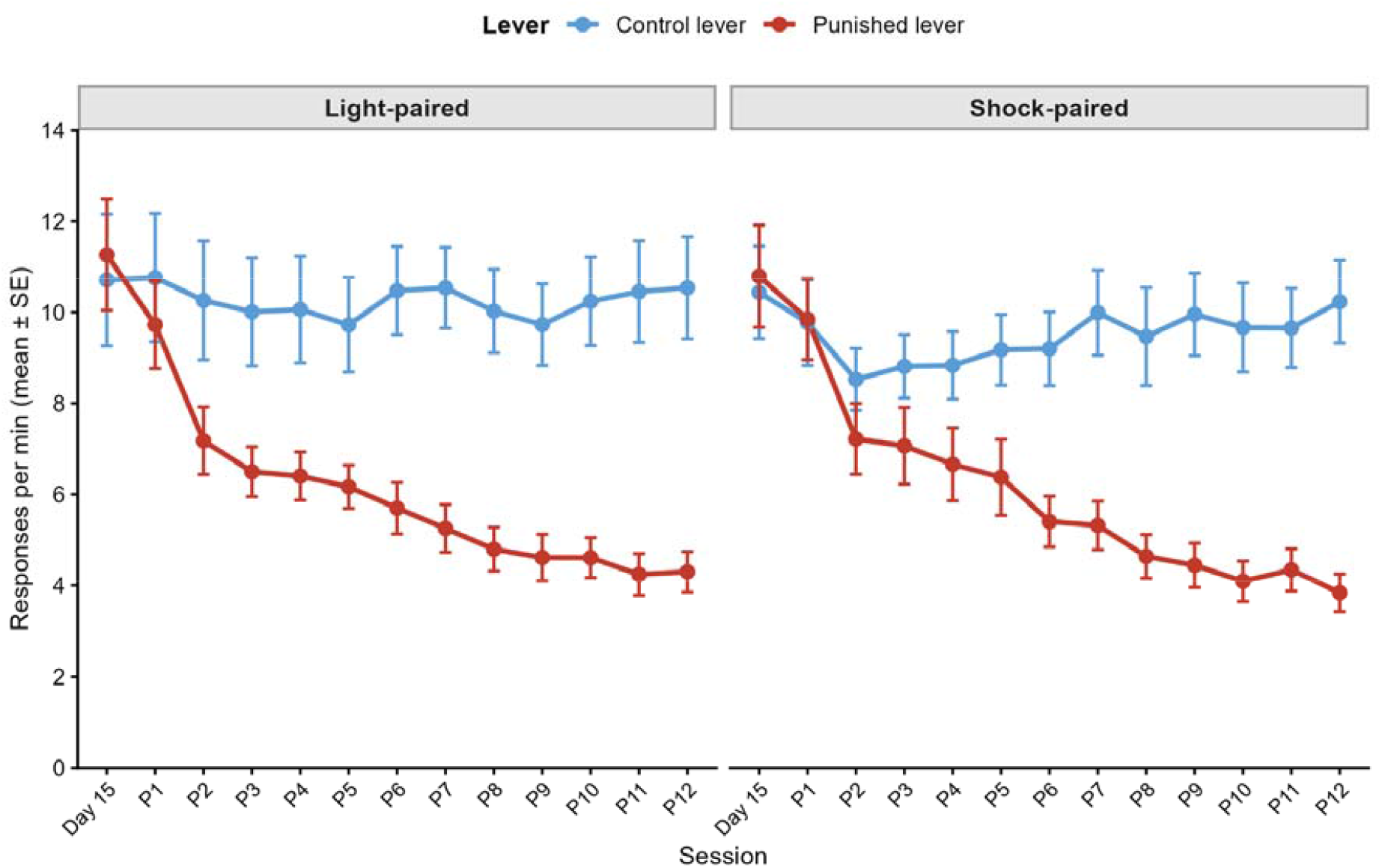
Lever press responding during punishment training. Responses per minute (mean ±SE) on control and punished levers during punishment training. Both groups showed equivalent baseline responding and progressive punishment-induced suppression, confirming group equivalence prior to counterconditioning (all Group effects: *ps* > .60). Error bars represent within-subjects SE.

Critically, all effects involving Group were nonsignificant: main effect of Group, F(1,44) = 0.27, *p* = .604; Group x Condition, F(1,44) = 0.24, *p* = .630; Group x Session, F(12,528) = 0.26, *p* = .994; Group x Condition x Session, F(12,528) = 0.67, *p* = .780. These null effects confirm that the shock-paired and light-paired groups were equivalent in baseline responding, punishment-induced suppression, and suppression dynamics throughout the punishment phase. Batch effects were also nonsignificant (all *ps* > .07). Thus, any subsequent group differences in outcome revaluation tests can be attributed to counterconditioning effects rather than pre-existing differences.

### Counterconditioning

During counterconditioning, one group received shock paired with food delivery (Shock-paired) while the control group received light-off paired with food (Light-paired). Figure 2A shows progressive acquisition of conditioned approach to the paired stimulus, with stronger conditioning in the Shock-paired group, suggesting greater shock salience. A Group x Stimulus x Session x Batch mixed ANOVA on difference scores (post-pre) revealed significant main effects of Group, F(1, 44) = 7.36, *p* = .009, *ges* = .035, Stimulus, F(1, 44) = 122.77, *p* < .001, *ges* = .363, and Session, F(7, 308) = 30.38, *p* < .001, *ges* = .153. Critically, a Group x Stimulus interaction, F(1, 44) = 65.10, *p* < .001, *ges* = .232, indicated differential conditioning: the Shock-paired group exhibited significantly stronger approach to the paired signal (M = 1.70) compared to both the Light-paired signal (M = 0.74) and the Shock-paired unpaired signal (M = 0.02), both *ps* < .001, whereas the Light-paired group showed more modest differentiation between paired (M = 0.74) and unpaired (M = 0.48) signals, *p* < .001. Although shock may elicit some unconditioned magazine approach in the Shock-paired group, the critical evidence for counterconditioning is within-group discrimination between paired and unpaired stimuli, which was significant in both groups.

**Figure 2.**
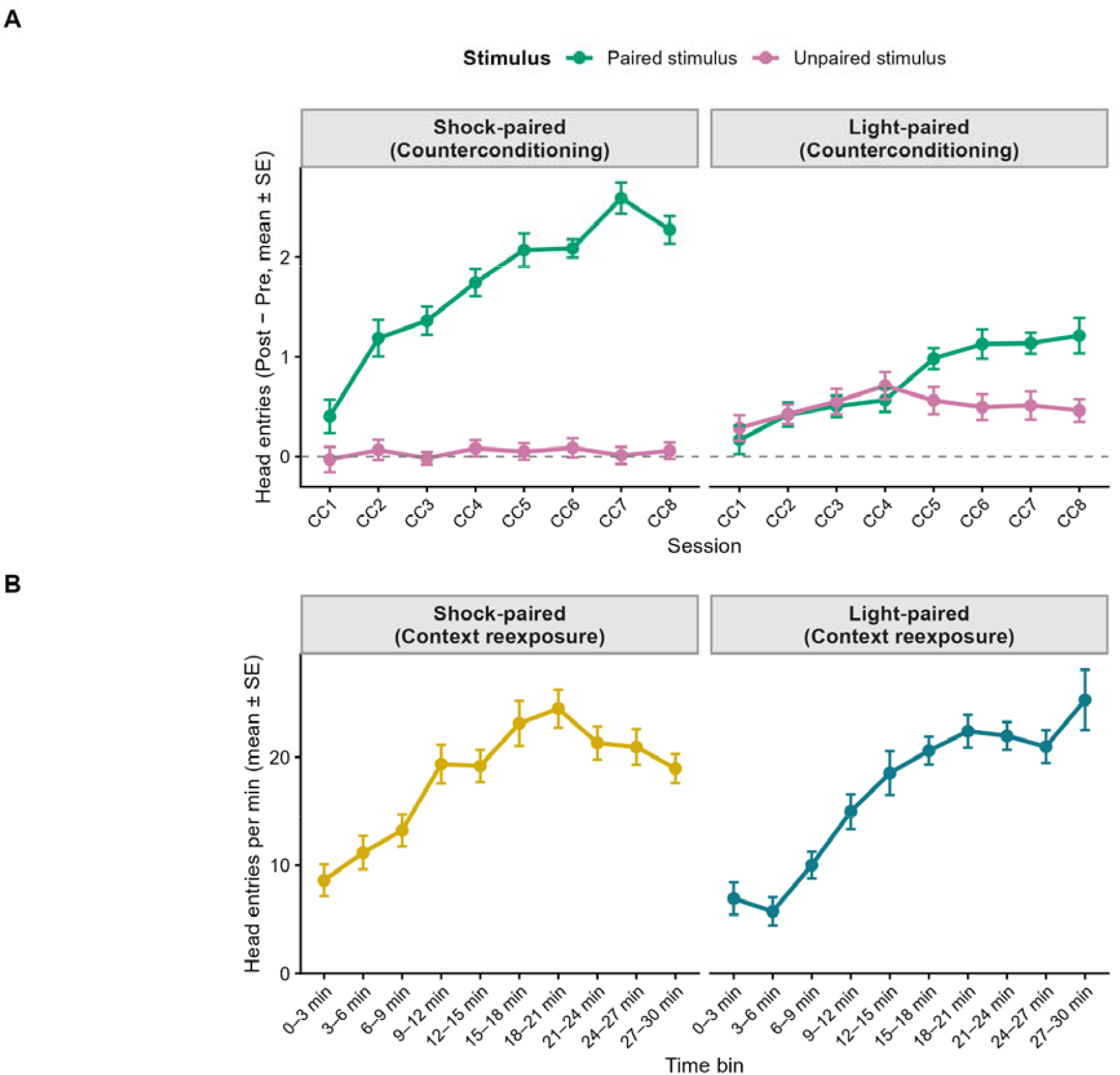
Appetitive counterconditioning and context reexposure. *Panel A.* Differential magazine approach (mean ± SE) during counterconditioning sessions. Both groups acquired a conditioned approach to the paired stimulus, with stronger conditioning in the shock-paired group, consistent with greater shock salience. *Panel B*. Head entries per minute (mean ± SE) during the context reexposure session. Both groups showed progressive increases in magazine approach across the session, with no significant group differences. Error bars represent within-subjects SE.

A significant Group x Stimulus x Session interaction, F(7, 308) = 3.45, *p* = .001, *ges* = .024, and a four-way Group x Batch x Stimulus x Session interaction, F(7, 308) = 2.31, *p* = .026, *ges* = .016, suggested varying acquisition dynamics. No main effect of Batch emerged, F(1, 44) = 2.17, *p* = .148. Notably, when the analysis was restricted to the last three sessions, the four-way interaction was no longer significant, F(2, 88) = 1.24, *p* = .293, *ges* = .003, indicating stabilized batch effects by training end. These results demonstrate successful appetitive conditioning of the shock signal, establishing a foundation for subsequent outcome revaluation effects.

### Revaluation test

Following counterconditioning, rats were re-exposed to the training context for 30 min with food on a RT 60-s schedule. A Group × Bin × Batch mixed ANOVA revealed a significant main effect of Bin, F(9, 396) = 25.27, *p* < .001, *ges* = .202, and a significant Group × Bin interaction, F(9, 396) = 1.93, *p* = .046, *ges* = .019, with no individual bin surviving Bonferroni correction (all *ps* > .06). Descriptively, the shock-paired group responded more early in the session while the light-paired group responded more at the end (see Figure 2B). These results support the interpretation of the revaluation test findings as reflecting a change in the aversive value of the shock rather than differential contextual conditioning.

Figure 3A shows that responding on both levers was initially low following extended exposure to the chambers without levers during counterconditioning, then progressively increased across the session. Critically, counterconditioning in the shock-paired group completely abolished the punishment persistence observed in the light-paired group, demonstrating the key revaluation effect (see Figure 3B).

**Figure 3.**
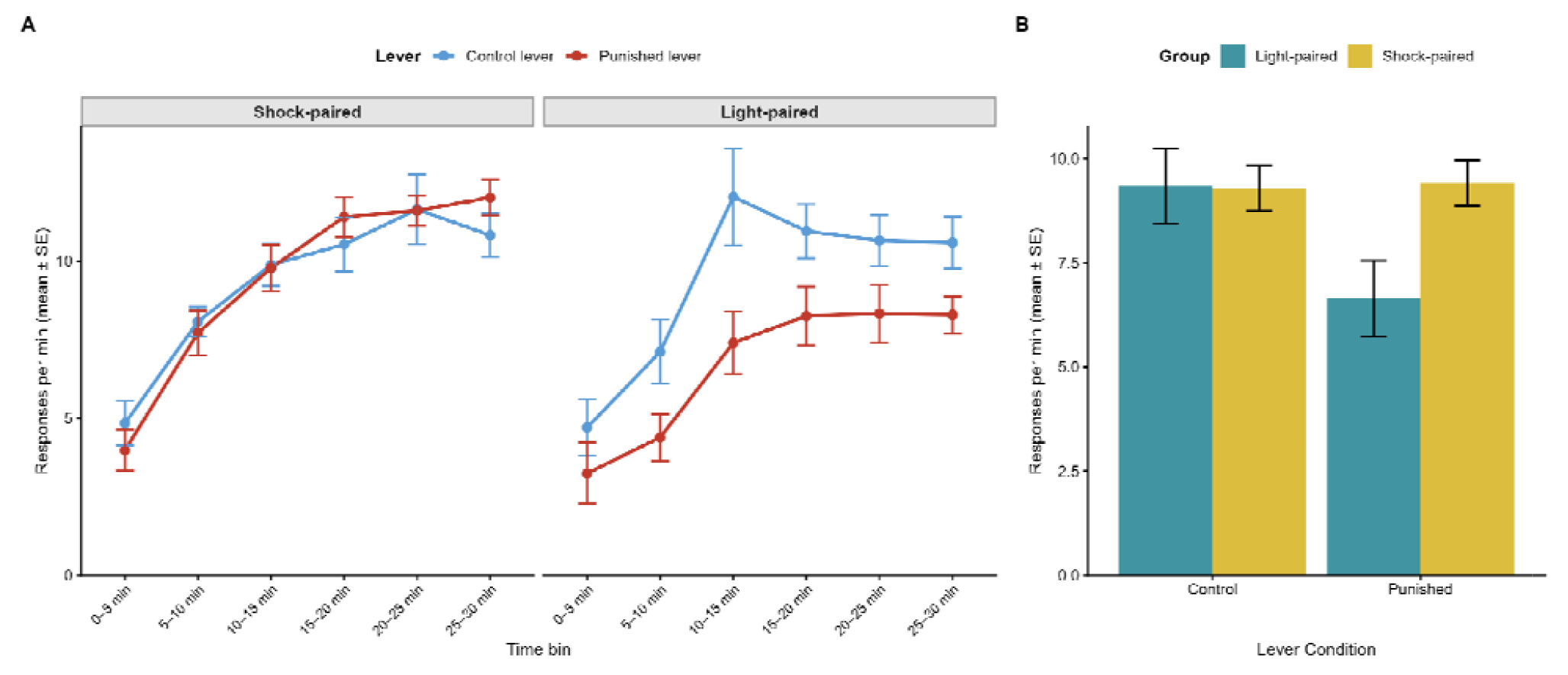
Lever-press responding during the Revaluation test. *Panel A.* Lever pressing (mean ± SE) across 5-min time bins during the Revaluation test. The shock-paired group showed complete recovery on the punished lever, while the light-paired group maintained suppression, demonstrating selective outcome revaluation. *Panel B*. Mean lever pressing aggregated across the full session. The shock-paired group responded equivalently on both levers, whereas the light-paired group responded significantly more on the control than the punishe lever. Error bars represent within-subjects SE.

Batch was not included as a between-subjects factor given that batch effects were nonsignificant across preceding phases. A Group x Condition x Phase mixed ANOVA revealed a significant three-way interaction, F(1,46) = 6.38, *p* = .015, *ges* = .009, driven by a robust Condition x Phase effect, F(1,46) = 73.49, *p* < .001, *ges* = .090, reflecting overall recovery on the punished lever, and a Group x Phase interaction, F(1,46) = 4.84, *p* = .033, *ges* = .012.

Planned comparisons within the Revaluation test showed the light-paired group responded significantly more on the control lever (M = 9.3 rpm) than the punished lever (M = 6.7 rpm), t(46) = 2.54, *p* = .015, *d* = 0.51, whereas the shock-paired group responded equivalently on both levers (control: M = 9.3 rpm; punished: M = 9.4 rpm), t(46) = -0.11, *p* = .911. Between groups, punished lever responding was higher in the shock-paired (M = 9.4 rpm) versus light-paired group (M = 6.7 rpm), t(85.6) = 2.22, *p* = .029, *d* = 0.64, with no difference on the control lever, p = .971. Recovery scores confirmed greater selective recovery in the shock-paired group (M = 6.52 rpm) than the light-paired group (M = 3.55 rpm), t(39.2) = 2.53, *p* = .016, *d* = 0.73. The linear mixed-effects model confirmed a significant Group x Lever interaction, t(46) = -2.45, *p* = .018, robust across temporal bins.

## Discussion

We tested whether instrumental punishment is controlled by the current value of the punisher. Rats were trained on a concurrent instrumental procedure in which both actions produced the same appetitive reinforcer, but one additionally produced shock. This within-subject design controls for generalized motivational effects, expressing suppression as a punished-versus-control contrast. We then manipulated the punisher’s value through counterconditioning, arranged so that both groups received equivalent shock and food exposure, but only the shock-paired group received explicit pairings of the appetitive reinforcer with the shock itself. Finally, instrumental performance was assessed in extinction with both actions available, but no shock delivered.

The key result shows that revaluation of the aversive outcome reliably reduced punishment suppression. The light-paired group showed the expected suppression of the punished relative to the control action, whereas in the shock-paired group the punished action fully recovered, becoming statistically indistinguishable from the control action. This pattern is exactly what a value-driven account predicts: changing the punisher’s value changes punished responding even in its absence, indicating that animals use an updated representation of the aversive outcome to motivate instrumental action.

Other procedures that generate response reduction do not necessarily involve the same type of representations. An omission schedule is a negative instrumental contingency in which performing a response causes the omission of an otherwise freely delivered outcome. Like punishment, omission contingencies suppress responding: animals learn to withhold the response to receive the outcome. Dickinson and colleagues (1998) trained rats to press two levers concurrently for food pellets while sucrose was delivered independently of responding. Pressing one lever postponed sucrose delivery, whereas pressing the other had no effect. This negative contingency suppressed responding on the omission lever relative to the control lever. Critically, when sucrose was subsequently devalued by pairing it with lithium chloride, devaluation had no effect on suppression: the omission effect was identical in devalued and non-devalued groups. This insensitivity suggests that suppression under omission is not mediated by a representation of the outcome’s current value, but rather by an inhibitory S-R association impervious to changes in outcome value (Colwill, 1991; Konorski, 1967; Rescorla, 1993).

The variation in the susceptibility of response suppression produced by punishment and omission contingencies to outcome revaluation parallels that observed with rewarded responding. In the case of rewarded behavior this variation has generated various dual-system theories. Psychological theories appeal to a goal-directed system paired with a habit system, which assumes that the rewarding outcome directly reinforces the response through the strengthening of a stimulus-response (S-R) association. Computational reinforcement theory draws a parallel distinction between model-based and model-free systems. The latter endows responses with the summed utilities of all states that have followed the response without retaining information about outcome content (Daw et al., 2005; Sutton & Barto, 1981). It is the model-free and habit system that explain responding impervious to outcome revaluation (Balleine & Dezfouli, 2019; Daw et al., 2005; Dickinson, 1994; Dolan & Dayan, 2013; Perez & Dickinson, 2020). In this approach, insensitivity to punisher revaluation would suggest that suppression is driven by inhibitory S-R learning rather than by a representation of the punisher’s current value.

The distinction between value-driven and habitual control of behavior readily explains why punishment fails to modify behavior under some conditions. For example, severe drug use persists in spite of severe and long-lasting aversive consequences. Evidence from both human and animal models suggest that persistent drug seeking transits from being outcome-value driven or goal-directed to habitual where the consequences are no longer being considered by drug users (Everitt & Robbins, 2016). Rats will maintain reward-seeking behavior even when presented with fear-related cues after extended cocaine exposure and consumption (Vanderschuren & Everitt, 2004), while humans with cocaine dependence will show impaired suppression of behavior in response to negative feedback compared to healthy controls (Ersche et al., 2016). Our results support the idea that punishment is a value-driven process that becomes habitual or value-insensitive under some environmental and psychological factors.

This habit view of punishment would seek to explain our results by an S-R inhibition account. However, it is unlikely that S-R inhibition could explain our results. As our data demonstrate, the punishment procedure did not generate any S-R inhibition in that the revaluation of the footshock totally abolished the difference between control and punished responding so that there was no residual suppression to be explained by S-R inhibition (see Figure 3). Furthermore, it is also unlikely that a classic account of punishment in terms of response competition (see Mackintosh, 1983, for review) was involved. The most obvious source of response competition would have been responding on the control lever, but this responding was similar in the revalued, shock-paired group and the non-revalued light-paired group.

There are, however, potential alternative accounts for our data. One of them, recently put forward by Crimmins and colleagues (2022) for reward learning, would argue that counterconditioning may reduce the aversive motivational properties of the location the outcome was delivered during training, rather than specifically updating the instrumental response-punisher representation. Under this account, recovery of responding on the punished lever would reflect a reduction in location-specific contextual fear rather than a genuine update of the causal representation. The concurrent within-subject design, while controlling for generalized motivational effects of training, does not fully resolve this alternative. A stronger test would employ a bidirectional lever procedure, in which the spatial position of the punished lever is counterbalanced across subjects, dissociating the instrumental response from its location.

This paper provides the first evidence that punishment actions can be governed by outcome value: animals update the value of an aversive outcome and integrate that updated value with response–outcome knowledge to regulate instrumental behavior. The broader implications are practical as well as theoretical: when control of punishment is value-driven, interventions that change the value of the punisher (or the inferred consequences of punished actions) should have principled, predictable effects on behavior—an assumption that had been so far difficult to justify empirically.

## Declarations

### Ethics approval

All experimental procedures involving animals were approved by the Institutional Animal Care and Use Committee at the authors’ institution and conducted in accordance with institutional guidelines for the care and use of laboratory animals.

### Consent to participate

Not applicable.

### Consent to publication

Not applicable.

### Availability of data and materials

The dataset and analysis code that support the findings of this study will be deposited in a public repository (e.g., OSF/Zenodo). Access links will be provided upon acceptance and will be included in the published article.

### Code availability

See “Availability of data and materials” statement above.

### Authors’ contributions

Author contributions have been masked for blind review.

## Acknowledgments

Acknowledgments information has been masked for blind review.

## Open Practices Statement

The dataset and analysis code supporting the findings of this study will be deposited in a public repository (OSF/Zenodo) upon acceptance; access links will be included in the published article. The experiment was not preregistered.

